# Differential effects of statins on the anti-dyskinetic activity of sub-anesthetic ketamine

**DOI:** 10.1101/2024.11.26.625570

**Authors:** Mitchell J. Bartlett, Carolyn J. Stopera, Stephen L. Cowen, Scott J. Sherman, Torsten Falk

## Abstract

Sub-anesthetic ketamine has been demonstrated to reduce abnormal involuntary movements (AIMs) in preclinical models of L-DOPA-induced dyskinesia (LID) and retrospective Parkinson’s disease case reports. In this study, we examined the effects on L-DOPA-induced dyskinesia of two statins alone and in combination with ketamine in unilateral 6-hydroxydopamine-lesioned male rats, the standard preclinical LID model. Sub-anesthetic ketamine attenuated the development of AIMs, while lovastatin only showed anti-dyskinetic activity at the beginning of the priming but did not prevent the development of LID. The polar pravastatin blocked the long-term anti-dyskinetic effects of ketamine, while the non-polar lovastatin did not. This study shows different classes of statins affect LID differentially, points to an important drug interaction and further supports ongoing clinical testing of sub-anesthetic ketamine to treat LID in individuals with Parkinson’s disease.

## 1. Introduction

The number of individuals with Parkinson’s disease (PD) is projected to reach 12 million by 2040.^1^ With growing access to levodopa (L-DOPA), the most effective therapeutic, a parallel epidemic of motor fluctuations including L-DOPA-induced dyskinesia (LID) is set to occur.^2^ The abnormal involuntary movements (AIMs) during LID can be just as debilitating as PD itself, and after 10-15 years of L-DOPA treatment more than 90% of individuals with PD will experience LID. However, pharmacotherapeutic treatments are limited; extended-release amantadine is the only FDA-approved drug, and has limited tolerability and efficacy in many individuals. Therefore, there is a great need for novel anti-dyskinetic pharmacotherapies. Our lab has rigorously evaluated sub-anesthetic ketamine in preclinical LID models.^3–7^ Ketamine is a multifunctional ligand, that includes *N*-methyl-D-aspartate receptor (NMDAR) antagonist and opioid receptor agonist activity,^7–9^ is FDA-approved and has a well-established safety profile.^10^ We previously showed that sub-anesthetic ketamine reduced AIMs in established LID and attenuated their development in preclinical models and improved or eliminated LID clinically in a retrospective chart review of five cases.^3,4,11^

We established that the long-term effects of ketamine depend upon brain-derived neurotrophic factor (BDNF)-signaling since the TrkB receptor antagonist, ANA-12, blocked ketamine from attenuating the development of AIMs.^4^ A recent study demonstrated that pravastatin, a polar 3-hydroxy-3-metylglutaryl-coenzyme A (HMG-CoA) reductase inhibitor, blocked the long-term antidepressive activity of ketamine by interfering with the direct binding of ketamine to the TrkB receptor.^12^ This suggests that cholesterol synthesis is required for the effects of ketamine on TrkB signaling and that a similar mechanism modifying BDNF-signaling could also be operative in the transduction of the anti-dyskinetic properties.^12^ In this study, we investigated the individual effects of two HMG-CoA reductase inhibitors, pravastatin (polar) versus lovastatin (non-polar), on the attenuation of AIMs alone and in combination with sub-anesthetic ketamine in a preclinical model of LID development.^4,6^

## 2. Methods

### 2.1. Animals

Male, Sprague-Dawley rats (10-weeks) were housed as published.^4^ Animal experiments were approved by The University of Arizona’s IACUC, in accordance with NIH and the ARRIVE 2.0 guidelines.

### 2.2. Unilateral 6-hydroxydopamine (6-OHDA) PD model

6-OHDA was injected unilaterally in the medial forebrain bundle (MFB) as published.^4^

### 2.2. Amphetamine-induced rotation test

Dopamine depletion following 6-OHDA was verified with amphetamine-induced rotations as published.^4^

### 2.3. Test drugs preparation

All drugs were compounded with vehicle, unless noted, and administered at 1 mL/kg via *intraperitoneal* (*i*.*p*.) or *subcutaneous* (*s*.*c*.) injections. Vehicle (0.9% USP-grade sterile saline, *s*.*c*.; VetOne, Boise, ID), ketamine (20 mg/kg, *i*.*p*.; VetOne), pravastatin (10mg/kg, *s*.*c*.; Cayman Chemicals, Ann Arbor, MI), lovastatin (10 mg/kg, *i*.*p*.; Cayman Chemicals), and L-DOPA (6 mg/kg, *i*.*p*.; MilliporeSigma, St. Louis, MO) combined with benserazide (12 mg/kg, *i*.*p*.; MilliporeSigma). Lovastatin was compounded as published.^13^

### 2.4. LID rating scale

LID severity was scored by a blinded investigator using the limb, axial, and orolingual (LAO)-AIMs rating scale as published.^3–7,14,15^

### 2.5. Treatment groups and experimental design

PD rats with a net ipsilateral rotation score ≥3 were distributed to six treatment groups: 1) Vehicle (V); 2) Ketamine (K); 3) Ketamine + Pravastatin (K+P); 4) Ketamine + Lovastatin (K+L); 5) Pravastatin (P); 6) Lovastatin (L). Statins or vehicle were administered for 14 days prior to first L-DOPA exposure on Day 0 and 16 days following the first L-DOPA exposure during AIMs development (**Fig. 1A**). Statins were injected daily 1-hour prior to peak HMG-CoA reductase activity, occurring 6-hours after the start of the dark cycle in the rat liver^16,17^ and brain.^18^ Ketamine or vehicle treatment: on Days 0, 7, and 16 using our established 10-hour ketamine paradigm (**Fig. 1B**).^3–6,19^ One animal was removed for non-specific health reasons.

**Figure 1.**
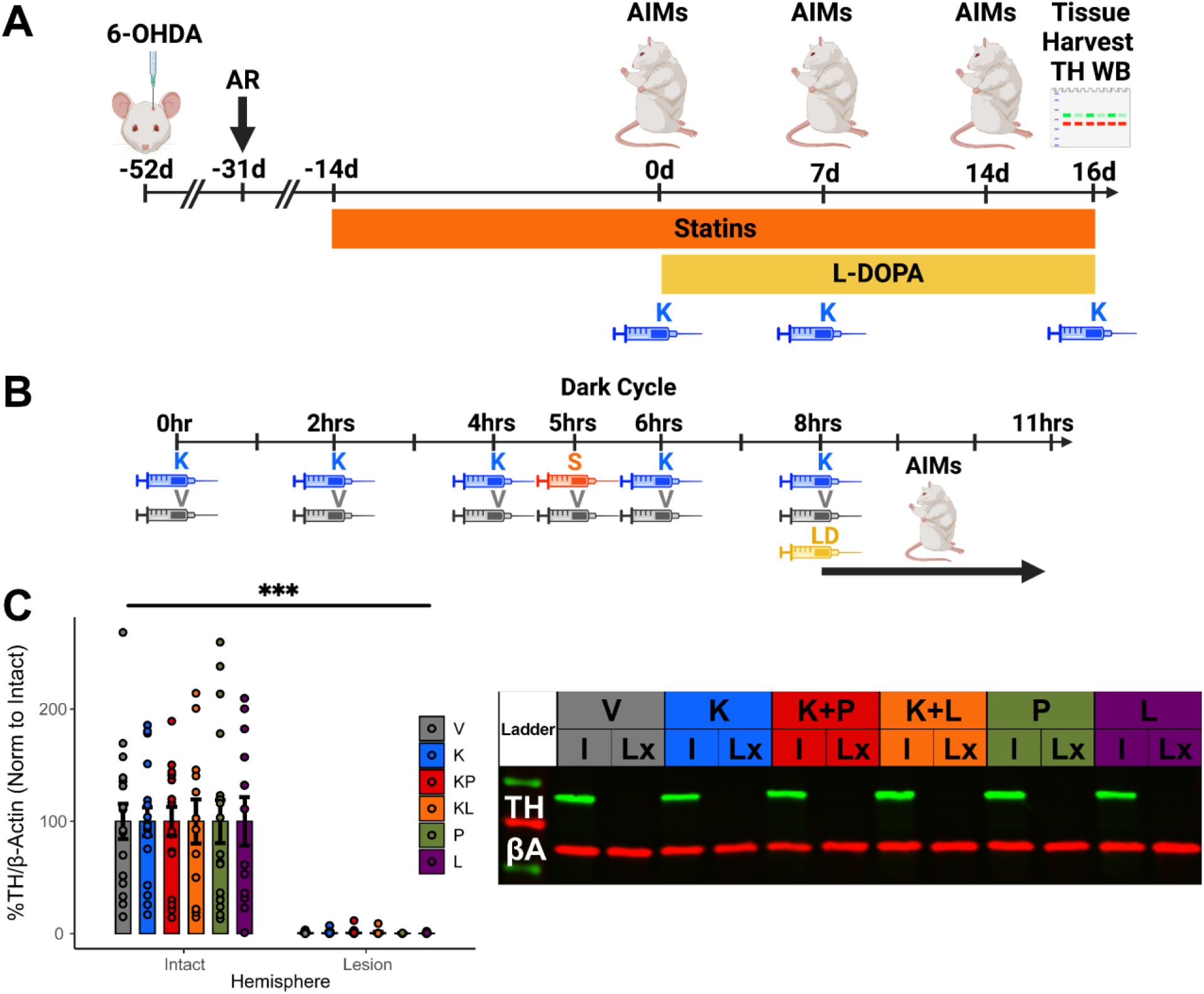
(**A**) Scheme of the preclinical PD model, statin/L-DOPA injection protocols, AIMs testing, and tissue harvest. (**B**) Scheme of the injection paradigm on AIMs testing days. K (ketamine); S (statins); V (vehicle); LD (L-DOPA). K was given on Days 0 and 7, but not Day 14 to test its long-term effect on LAO-AIMs. (**C**) Verification of 6-OHDA lesion in striatal tissue. Plotted as % loss (mean ± SEM) in the intact vs. lesion hemisphere. Two-way ANOVA, with Bonferroni *post hoc* tests, ***p<0.001, n = 12-17 per group. V (vehicle, grey); K (blue); K+P (K + pravastatin, red); K+L (K + lovastatin, orange); P (pravastatin, green); L (lovastatin, purple). Inset shows example blot. βA (Beta-actin); TH (tyrosine hydroxylase); I (intact); Lx (lesion). Figures 1A (CJ2763GY9I) and 1B (HZ2763HC8C) were created with BioRender.

### 2.6. Tissue preparation and western blot analysis

Striatal tissue was stained for tyrosine hydroxylase (TH) and analyzed to confirm dopamine depletion as published.^7^

### 2.7. Statistical analysis

Western blot analysis was performed with a two-way ANOVA, with Bonferroni *post hoc* tests using GraphPad Prism 10.0 software (GraphPad Software, La Jolla, CA). Analyses of LAO-AIMs were performed using Kruskal-Wallis tests, with Dunn’s multiple comparisons and Holms *post hoc* corrections tests, using R 4.4.1 (R Core Team, 2024). The null hypothesis was rejected when p<0.05. All values are represented as mean ± SEM.

## 3. Results

### 3.1 Verification of the hemi-parkinsonian lesions

Unilateral microinjection of 6-hydroxydopamine (6-OHDA) in the MFB results in a >90% reduction in striatal dopamine. Average amphetamine-induced rotations per minute (mean ± SEM) were: 1) vehicle (6.7 ± 0.46); 2) ketamine (6.6 ± 0.49); 3) ketamine + pravastatin (6.4 ± 0.43); 4) ketamine + lovastatin (6.1 ± 0.75); 5) pravastatin (6.4 ± 0.41); and 6) lovastatin (6.3 ± 0.63). Depletion of striatal dopaminergic terminals (F[1,172] = 204, p<0.001, two-way ANOVA, Bonferroni *post hoc* tests) was verified using post-mortem semi-quantitative western analysis of TH (**Fig. 1C**).

### 3.2. Effect of Ketamine and Statins on LAO-AIMs

We treated unilateral 6-OHDA lesioned PD rats with statins for two weeks prior to their first exposure to ketamine or vehicle, plus L-DOPA (**Fig. 1A**). On Days 0 and 7 rats were exposed to our previously established 10-hour ketamine treatment paradigm^3,4,6,19^ plus statins (**Fig. 1B**). On Day 0, with first exposure to L-DOPA, ketamine alone (p=0.0432) attenuated LAO-AIMs by 58% (**Fig. 2A**), while ketamine + lovastatin (p=0.0104) further attenuated LAO-AIMs by 77%, when both were compared to vehicle (**Fig. 2B**). Alone, lovastatin did not significantly reduce LAO-AIMs; however, pravastatin significantly increased LAO-AIMs when compared to lovastatin by 121% (p=0.040) and a trend of a 90% increase compared to vehicle did not reach statistical significance (**Fig. 2C**). On day 7 ketamine reduced LAO-AIMs by 54% (p=0.0049), but when combined with pravastatin, had no significant effect compared to vehicle (**Fig. 2D**), whereas ketamine combined with lovastatin significantly decreased LAO-AIMs by 75% (p=0.0005) compared to vehicle (**Fig. 2E**). In the absence of ketamine, lovastatin alone significantly reduces LAO-AIMs by 48% (p=0.0315) and 55% (p=0.0032) as compared to both vehicle and pravastatin treated groups, respectively. There is no difference between the vehicle and pravastatin treated groups by Day 7 (**Fig. 2F**). On Day 14, to test for ketamine’s long-term effects, LAO-AIMs in the group treated with ketamine alone were significantly reduced by 48% (p=0.0376) compared to the vehicle treated group and by 43% (p=0.0376) compared to the ketamine + pravastatin treated group (**Fig. 2G**). Ketamine + lovastatin significantly reduced LAO-AIMs by 62% (p=0.0049) compared to vehicle (**Fig. 2H**). By Day 14 there was no significant difference between lovastatin or pravastatin and vehicle (**Fig. 2I**).

**Figure 2.**
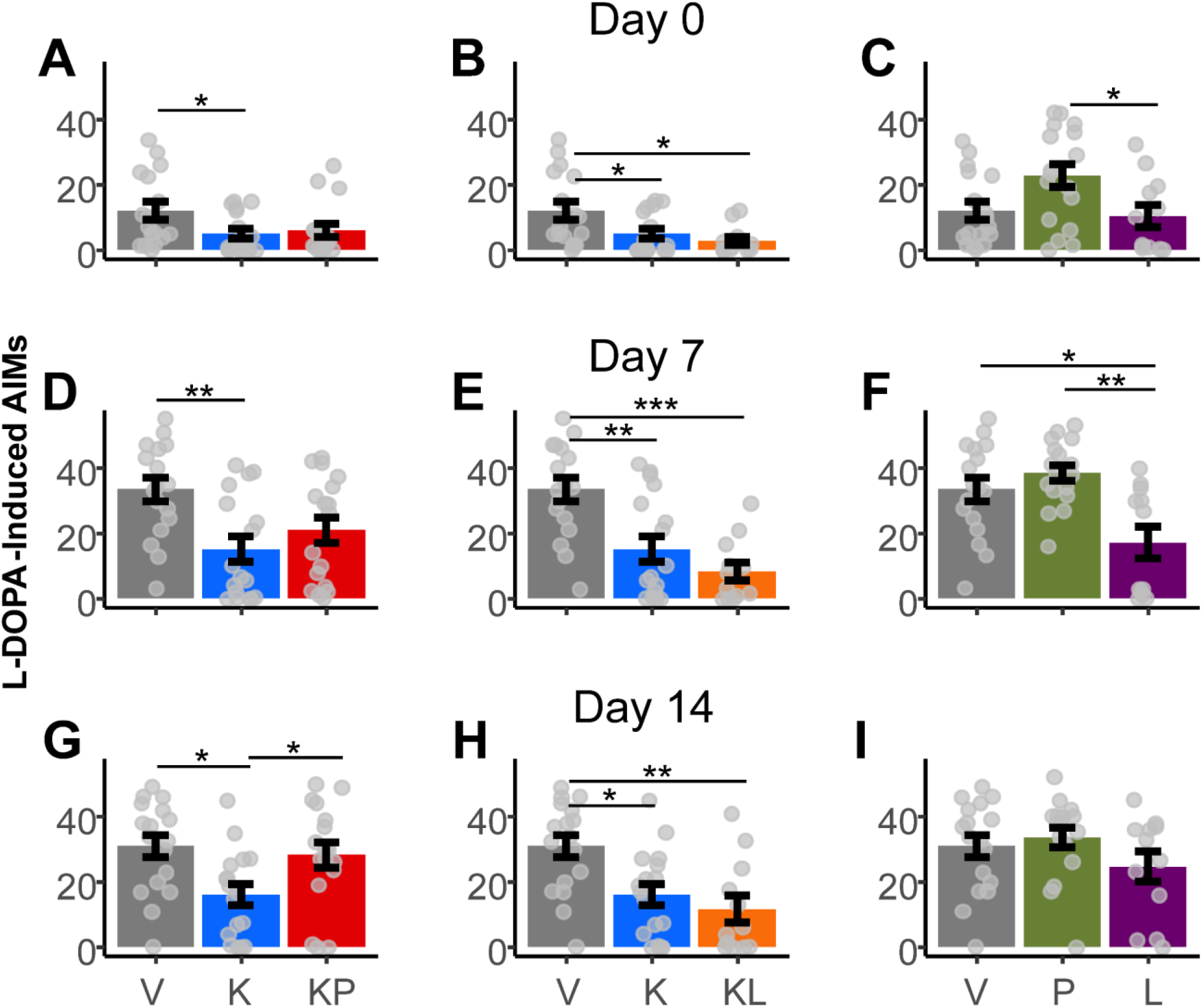
Pravastatin but not lovastatin blocks the long-term anti-dyskinetic activity of ketamine. (**A**) The LAO-AIMs (mean ± SEM) showed an acute reduction by ketamine (K). (**B**) K and ketamine + lovastatin (K+L) significantly reduced LAO-AIMs compared to vehicle (V) treated rats on Day 0. (**C**) Pravastatin (P) significantly increased LAO-AIMs compared to Lovastatin (L) and shows a trend vs. Vehicle, suggesting a possible sensitization effect at first exposure to L-DOPA. (**D**) Ketamine significantly reduced LAO- AIMs on Day 7. (**E**) K+L significantly reduced LAO-AIMs at Day 7. (**F**) Lovastatin alone showed anti-dyskinetic activity by Day 7, significantly reducing LAO-AIMs compared to both vehicle and pravastatin. (**G**) K+P has no long-term anti-dyskinetic effects suggesting that pravastatin interferes with the long-term anti-dyskinetic activity of K seen 7-days after the last ketamine treatment. (**H**) K and K+L continue to demonstrate long-term anti- dyskinetic activity at Day 14. (**I**) By Day 14 there is no significant improvement in LAO- AIMs by lovastatin alone, demonstrating that the anti-dyskinetic activity of lovastatin is a short-term effect and does not prevent LID development. Kruskal-Wallis with Dunn’s tests and Holms *post hoc* correction, *p<0.05, **p<0.01, ***p<0.001; n = 12-17 per group.

## 4. Discussion

The role of NMDAR-mediated signaling in LID is well-established^20^ resulting in the FDA approval of extended-release amantadine to treat LID. However, amantadine is a weak NMDAR antagonist with limited efficacy in reducing LID in individuals with advanced PD and poorly tolerated due to hallucinosis, in addition to other central nervous system (CNS) side-effects.^21^ Therefore, there is an urgent need for better treatments. Sub-anesthetic ketamine could fulfill this need. Ketamine is a multi-ligand drug, with improved NMDAR binding affinity compared to amantadine, and FDA approved for treatment-resistant depression.^22^ Preclinical studies^3–7^ and a retrospective case report^11^ have identified sub-anesthetic ketamine as an excellent candidate drug to treat LID. We recently completed a Phase 1 open-label clinical trial of 10-hour sub-anesthetic ketamine infusions in subjects with refractory LID (NCT06021756), showing safety, tolerability and early indications of long-term efficacy to support a planned placebo-controlled Phase 2B clinical trial.

We previously demonstrated that blocking BDNF from binding its TrkB receptor prevented the long-term anti-dyskinetic effect of ketamine,^4^ providing evidence that, in addition to acute NMDAR-antagonism and opioid-agonism, LID prevention by ketamine is also mediated by TrkB receptor activation and downstream effectors in mTOR signaling. These data agree with studies showing that ketamine-treatment leads to increased BDNF and mTOR expression in models of depression.^23^ Conversely, these antidepressant effects are blocked in BDNF null mutant and Trk dominant-negative transgenic mice.^24^ Therefore, mounting evidence suggest that ketamine exerts its potent antidepressant effect via a BDNF-mediated transduction pathway. We hypothesized that a similar mechanism underlies the potent anti-dyskinetic effect of ketamine.

In this study, we tested the ability of ketamine, when combined with a polar (pravastatin) or a non-polar (lovastatin) statin drug, to reduce LID in a preclinical rodent model. First, we confirmed previous results that ketamine alone significantly attenuated the development of AIMs in parkinsonian rats treated with L-DOPA. We then evaluated the independent effects of each statin on AIMs, validating and extending a prior study by Bezard’s group^13^ demonstrating that, while lovastatin has some early anti-dyskinetic activity, it does not lead to the long-term attenuation of LID achieved by ketamine alone. Importantly, lovastatin does not interfere with the anti-dyskinetic activity of ketamine. In contrast, pravastatin interferes with ketamine binding to the TrkB receptor and blocks the long-term anti-dyskinetic effect of ketamine. This result is consistent with Casarotto et al., who show pravastatin interferes with the long-term antidepressive actions of ketamine.^12^ Their experiments demonstrated that the long-term antidepressive effect of ketamine involves its direct binding to the TrkB receptor, leading to increased membrane availability, TrkB receptor activation, thereby upregulating this pathway, and ultimately leading to neuroplastic changes in the CNS.^12^

The anti-dyskinetic activity of ketamine has been only evaluated and verified in male rats, and given the mechanistic nature of this study we excluded females. This limitation needs to be addressed in future studies, especially given the reported sex-specific effects of ketamine in models of schizophrenia and depression.^25^

As statins are regularly used by many individuals with PD, these data demonstrate the need to evaluate and consider their effects when planning Phase 2 and 3 clinical trials of sub-anesthetic ketamine for LID treatment. The use of polar statins should be considered as an exclusion criterium, and potential subjects switched to a non-polar statin. In addition, the unexpected trend of a possible sensitization after first exposure to pravastatin and L-DOPA raises concerns in general about the use of polar statins in individuals with PD and should be further evaluated. In conclusion, this study points to an important drug interaction and provides additional support for repurposing of sub-anesthetic ketamine for the treatment of LID in individuals with PD.

## CRediT authorship contribution statement

**Mitchell J. Bartlett:** Writing – original draft, Validation, Visualization, Methodology, Supervision, Investigation, Formal analysis. **Carolyn J. Stopera**: Writing – review & editing, Methodology, Investigation, Formal analysis. **Stephen L. Cowen:** Formal analysis, Writing – review & editing. **Scott J. Sherman:** Writing – review & editing, Funding acquisition. **Torsten Falk:** Writing – review & editing, Writing – original draft, Supervision, Resources, Funding acquisition, Design.

## Data availability

Data will be made available on reasonable request.

